# Why This Code? A Constrained Mapping Framework for the Evolutionary Stability of the Canonical Genetic Code

**DOI:** 10.64898/2026.08.28.747708

**Authors:** Rentian Lin, Chenxiao Wang, Yining Hu, Chuanrui Wang

## Abstract

The canonical genetic code is used by most known forms of life, yet explaining its historical origin and present-day functional performance requires comparison with the enormous space of possible codon-to-output assignments. Here, the code is formulated as a hierarchy of constrained mapping problems spanning codeword length, degeneracy composition, synonymous-block partitioning, semantic assignment and a coarse decoder layer. Exact structural analyses identify triplets as a Pareto choice under a fixed-length full-codebook model and show that anonymous degeneracy statistics alone do not explain the canonical profile. Within a fixed canonical block architecture and under specified objective functions, recurrent AAindex-based rule learning contracts the 20! amino-acid assignment space to 2.72 × 10^11^ admissible mappings, from which 10^8^ complete codes are sampled. In this screened conditional candidate library, the standard genetic code ranks in the best 0.9749% under the equal-weight three-objective score and in the best 1.801% when accessible replacement diversity is added. Sensitivity analyses show that this position is broad across many, but not all, tested objective weights and aggregation rules. These results describe a conditional multi-objective compromise; they do not establish global optimality, historical inevitability or cellular feasibility of decoder redesign.

## 1 Introduction

The standard genetic code is one of the most conserved features of living systems. With limited exceptions in mitochondria and some microbial lineages, nearly all organisms use the same triplet code to translate 64 codons into 20 amino acids and termination signals. This near universality suggests that the code was established early in evolution and subsequently maintained under strong constraints, yet the reasons why this particular codon-to-output mapping became fixed remain debated.

Crick’s frozen accident hypothesis argues that, once a coding scheme was adopted by early life, later changes would have caused widespread mistranslation and were therefore strongly selected against^1^. Error-minimization theories instead emphasize the internal organization of the code. Codons that are easily interconverted by point mutation or translational error often encode amino acids with similar physicochemical properties, reducing the functional cost of errors^2,3^. Other models focus on stereochemical affinity between amino acids and cognate codons or anticodons^4^, or on coevolution between the genetic code and amino acid biosynthetic pathways^5^. These views are not mutually exclusive, and each captures a plausible component of code evolution. However, none alone establishes whether the present code should be regarded as a historical contingency, an optimized solution, or a compromise shaped by multiple constraints^6^.

A central difficulty is the size and structure of the possible code space. If codon assignments are treated as arbitrary mappings from 64 codons to 20 amino acids and stop signals, the number of alternatives is enormous. Moreover, many random codes are biologically implausible because they ignore triplet reading, degeneracy, stop codons, codon-block organization and other structural features of translation. This makes it necessary to compare the standard genetic code not against unconstrained random mappings, but against alternative codes generated under comparable biological and combinatorial constraints.

Here, we formulate the genetic code as a constrained mapping problem from 64 codons to 20 amino acids and one termination signal. The search space is reduced by specifying codon length, degeneracy limits, degeneracy composition, codon-block partitioning and block-to-output assignment. To quantify amino-acid similarity during assignment-space screening, we use AAindex, a curated database of numerical indices describing the physicochemical and biochemical properties of amino acids^26^. Candidate codes are then evaluated using metrics related to mutational robustness, GC balance and encoding diversity, with *Escherichia coli* MG1655 protein-coding sequences used as a reference biological test set^7,8^. This framework allows the standard genetic code to be examined as a structured reference point within a biologically constrained space, rather than as an isolated historical outcome.

## 2 Methods

### 2.1 Hierarchical representation of genetic-code space

The nucleotide alphabet was Σ={A,C,G,U}. A genetic code was represented as Ω=(L,R,H,P,ψ), where L is code-word length, H=(h_1_,…,h_21_) is the multiset of output degeneracies satisfying Σ_*i*_h_*i*_=4^*L*^, R=max(H), P={B_1_,…,B_21_} is an anonymous partition of Σ^*L*^ with |B_*i*_|=h_*i*_, and ψ is a bijection from the 21 anonymous blocks to 20 amino acids plus termination. Analyses were conditional and layer specific: the L analysis used only code capacity and the quaternary Hamming graph; H/R analyses used integer compositions of 4^*L*^ into 21 positive parts; P analyses used codon membership and single-nucleotide adjacency without amino-acid labels; and ψ analyses introduced amino-acid semantics only after P had been fixed. This ordering prevents downstream biological information from being used to explain an upstream structural choice. For the principal large-scale experiments, L=3, the 20 canonical sense blocks and the UAA/UAG/UGA termination block were fixed, Stop was not permuted, and ψ ranged over the 20! bijections between the sense blocks and amino-acid labels^9–16^.

For the local third-position partition calculation, each of six canonical four-codon boxes containing a 2+2 synonymous split was treated independently. A third-position transition had weight *κ* and a transversion had weight 1. The cut weight induced by the canonical U/C | A/G split was compared exactly with the two alternative pairings. For the code-length calculation, vertices of the quaternary Hamming graph were all words in Σ^*L*^ and edges joined words differing at one position. Surjective 21-way partitions were evaluated by RatioCut; exact edge-isoperimetric profiles and subset dynamic programming were used to minimize the objective over permitted block-size compositions. These structural calculations were kept separate from the fixed-block ψ experiment reported below.

### 2.2 AAindex recurrent-rule screen and exact candidate generation

Let *W* _*ij*_ denote the number of undirected, single-nucleotide Hamming edges joining canonical sense blocks *B*_*i*_ and *B*_*j*_. Internal edges contribute zero, and edges incident to the fixed Stop block were excluded. Screening used a symmetric 20 × 20 AAindex distance matrix *D*_AAindex_, derived from physicochemical AAindex1 features and scaled to mean off-diagonal distance one^26^. No ProteinGym value, directional average or host-proteome quantity entered screening. For a complete semantic assignment ψ, the screening cost was

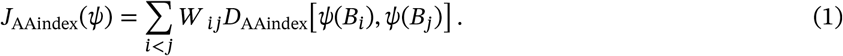

The matrix is symmetric by construction, so equation (1) is direction-neutral and does not assume a wild-type-to-mutant source distribution. Lower values place amino acids with smaller AAindex separation on adjacent codon blocks.

Independent uniform permutations were generated by a vectorized Fisher–Yates shuffle. Three independent seed groups were used, each containing three rounds of 10,000,000 complete mappings. In each round, the lowest-cost fraction *q* = 0.10 defined the discovery tail. For every block *i* and amino-acid label *a*, relative representation was

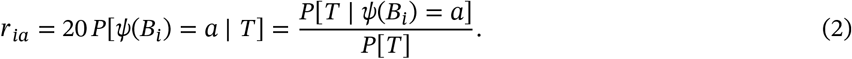

In equation (2), *r*_*ia*_ is the relative representation of assigning amino acid *a* to block *B*_*i*_ in discovery tail *T*; the factor 20 sets the uniform expectation to one. A noncanonical assignment was a provisional exclusion when *r*_*ia*_ < 1. A rule had to recur in at least two of three rounds within a seed group and in all three independent seed groups. Canonical assignments were protected. The reference screen yielded 196 recurrent exclusions, which were compiled into a 20 × 20 Boolean allowed-assignment matrix.

The number of admissible complete mappings was calculated exactly as the permanent of the allowed matrix using subset dynamic programming:

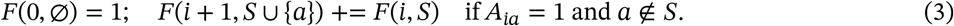

Here, *F*(*i, S*) is the number of partial bijections assigning the first *i* blocks to label subset *S*, and *A*_*ia*_ records whether label *a* is allowed for block *i*. The terminal value *F*(20, {1, …, 20}) gave 271,542,412,800 admissible mappings. The same completion-count table supported exact uniform generation: a rank drawn from [0, *F* − 1] was recursively unranked by subtracting legal-next-label completion counts. This generated 100,000,000 complete mappings without rejection sampling. The exclusions are recurrent empirical proposal constraints, not proofs that excluded mappings are biologically impossible.

### 2.3 Three-objective scoring: mutation robustness, encoding entropy and local GC stability

The principal landscape used three objectives calculated only after candidate generation. Mutation robustness used a frozen directional ProteinGym v1.3 wild-type-to-mutant penalty matrix *C*_dir_^17^. ProteinGym was not used to learn the AAindex screen. Under each candidate mapping, a source amino acid was drawn uniformly, a synonymous codon was drawn uniformly, and one of the nine single-nucleotide substitutions was drawn uniformly. The directional robustness cost was

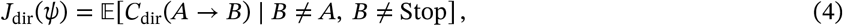

where *A* and *B* are the source and translated target amino acids. Lower expected directed missense cost was favourable. The exact assays, filtering, score orientation, aggregation and missing-value treatment used to derive this 20 × 20 matrix must be frozen with the analysis code; ProteinGym itself does not supply this project-specific matrix.

Encoding entropy used amino-acid occurrence frequencies pooled from the frozen *Escherichia coli* K-12 MG1655 reference proteome (UniProt proteome UP000000625; 4,403 records). This was an occurrence-weighted, not protein-abundance-weighted, reference. If *d*_*b*_ is the number of synonymous codons in fixed block *B*_*b*_ and *b*_ψ_(*a*) is the block assigned to amino acid *a*, synonymous codons were assumed to be equiprobable and the expected sequence-choice capacity was^18–19^

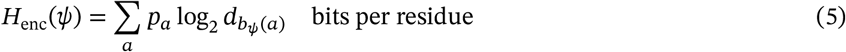

In equation (5), Henc(ψ) is the expected encoding entropy per residue under mapping ψ; a indexes amino-acid identities; p_*a*_ is the observed proteome frequency of amino acid a; bψ(a) identifies the fixed block assigned to a by ψ; and d_{bψ(a)} is that block’s number of synonymous codons. Thus, an occurrence of amino acid a contributes log_2_d bits when it is assigned to a block containing d synonymous codons, and the proteome-wide score is the amino-acid-frequency-weighted mean of these contributions. Higher Henc indicates a larger expected number of synonymous nucleotide sequences capable of encoding the same reference proteome under the uniform-codon assumption; it does not measure the realized entropy of native codon usage.

Local GC stability was computed analytically rather than by sampling reverse-translated sequences. For every fixed block b, μ_*b*_ and v_*b*_ were the mean and variance of the number of G/C nucleotides among its synonymous codons under uniform use. For each overlapping 20-residue window (60 nucleotides) with amino-acid count vector x, the expected squared deviation of window GC fraction G/(3w) from 0.5 was

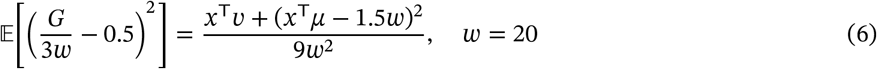

In equation (6), E denotes expectation over uniform synonymous-codon choices; G is the random number of G or C nucleotides in the window; w is the number of amino-acid residues per window and was fixed at 20; x is the 20-component amino-acid count vector of the window; μ and v are vectors containing, for the block assigned to each amino acid, the mean and variance of its per-codon G/C count; and the superscript T denotes vector transpose. All codon-aligned windows were included with a one-codon stride; windows were averaged equally within each eligible protein and proteins were then averaged equally. Lower GC mean-squared error was favourable. This score measures intrinsic local composition under a uniform synonymous model and does not fit candidates to native codon bias, transcription or ribonucleic acid (RNA) folding.

Raw objectives were placed on a common dimensionless scale using empirical midranks in the complete supplied 100-million-code library. For a lower-is-better objective with candidate value x, empirical loss was

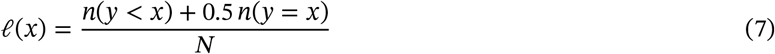

In equation (7), *l*(x) is the loss percentile of candidate value x; y denotes values of the same objective among the N supplied candidate codes; and n(·) counts candidates satisfying the stated relation. For a higher-is-better objective, y<x was replaced by y>x. Utility was then defined as

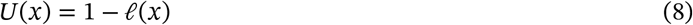

In equation (8), U(x) is the empirical utility of value x, with zero worst and one best in the supplied library. The equal-weight three-objective composite was

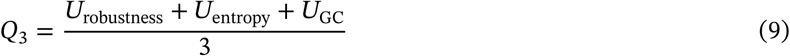

In equation (9), Q_3_ is the composite three-objective quality; Urobustness, Uentropy and UGC are the utilities of mutation robustness, encoding entropy and local GC stability, respectively. The SGC was evaluated explicitly against the same empirical distributions. For a candidate permutation π relative to the SGC permutation πSGC, the minimum label-transposition distance was

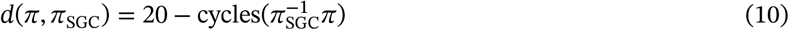

In equation (10), d is the minimum number of pairwise label exchanges; π is the candidate block-label permutation; πSGC is the SGC permutation; πSGC^−1^π is their relative permutation; and cycles(·) counts its disjoint cycles, including fixed points. At each integer distance d, enrichment of globally top-1% codes was

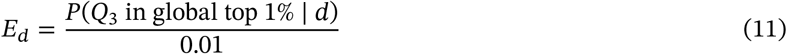

In equation (11), E_d is the fold enrichment at distance d and P denotes the within-distance empirical probability. Only strata with at least 1,000 sampled codes were displayed, and uncertainty was reported as a two-sided 95% Wilson interval for the within-stratum binomial proportion. Equal weighting was used as a transparent descriptive scalarization; Pareto comparisons were retained as a weight-free sensitivity analysis^20–21^.

### 2.4 Accessible replacement diversity and constrained decoder-unit redesign

Accessible replacement diversity was calculated for each fixed-block mapping as follows. A source amino acid was drawn uniformly, one of its synonymous codons was drawn uniformly, and one of the nine one-nucleotide substitutions was drawn uniformly. After global conditioning on a sense missense outcome, two independent target amino acids B_1_ and B_2_ were drawn from the conditional distribution for the same source. Their separation was scored with the unmodified directional ProteinGym-derived cost table:

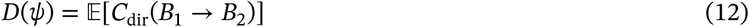

In equation (12), D(ψ) is accessible replacement diversity under mapping ψ; E denotes expectation over the declared source-codon-mutation sampling process conditional on a sense missense event; B_1_ and B_2_ are independent target amino acids drawn for the same source amino acid; and Cdir(B_1_→B_2_) is the directional ProteinGym-derived replacement cost from B_1_ to B_2_. Because B_1_ and B_2_ are identically distributed, ordered target pairs occur with equal probability in both orientations, so the expectation is insensitive to the antisymmetric component of the table even though the original directional entries were retained. Higher D indicates broader accessible outcomes in empirical replacement-cost space; it is not a measured probability of beneficial mutation. Four-objective quality was

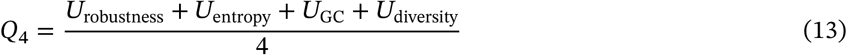

In equation (13), Q_4_ is the four-objective composite and Udiversity is the empirical utility of accessible replacement diversity; the other utilities are defined in equation (9). Q_4_ was analysed with the same empirical-rank procedure^17,22^.

Fixed-block relabeling leaves the raw number of distinct SNV target blocks invariant, so target-richness optimization was performed in a larger decoder-unit space. The 61 sense codons were grouped by SGC amino-acid identity and first-two-base box. Within each group, U/C and A/G third-position pairs were treated as atomic units when both codons were present; remaining codons were singletons. This coarse wobble model yielded 32 units (29 pairs and three singletons). It is a transparent mechanistic constraint, not a reconstruction of the complete modified-tRNA repertoire of MG1655^11,13–14,23^. AUG remained a singleton fixed to Met, the three Stops remained fixed, and proposals exchanged amino-acid identities only between decoder units of equal size, thereby preserving amino-acid degeneracies and preventing an atomic unit from being split.

For a candidate decoder state, accessible richness was

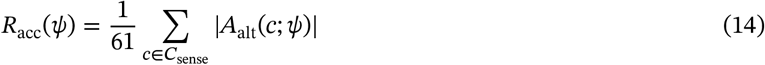

In equation (14), Racc(ψ) is mean accessible richness; Csense is the set of 61 sense codons; c is a source codon; Aalt(c;ψ) is the set of distinct alternative amino-acid labels reached from c through sense one-nucleotide neighbours under decoder state ψ; and |·| denotes set cardinality. Directional missense cost assigned equal prior weight to each source amino acid and equal weight to all directed sense-to-sense one-base edges originating from codons assigned to that source. The 60-nucleotide GC score was recalculated from the candidate’s dynamic synonymous sets using the same sufficient-statistic expression above. Feasibility required

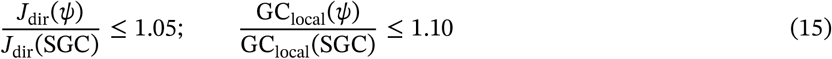

In equation (15), Jdir is the directional missense cost and GClocal is the 60-nucleotide local GC-stability cost; each ratio compares a candidate decoder state ψ with the SGC. Engineering burden was measured by the minimum number of nucleotide substitutions required to recode the MG1655 reference coding sequence (CDS) collection:

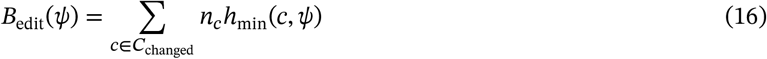

In equation (16), Bedit(ψ) is total recoding burden; Cchanged is the set of codon types whose amino-acid meaning changes; n_c is the genomic occurrence count of codon c in the reference CDS collection; and hmin(c,ψ) is the smallest nucleotide Hamming distance from c to any codon that retains c’s original amino-acid meaning under ψ.

All decoder-unit exchanges within change radius two were enumerated exactly. For radii 4, 6, 8, 10 and 12, simulated annealing used eight independent restarts of 200,000 proposal steps, with seed 20260803. The search score maximized richness while subtracting the minimum substitution fraction and large penalties for constraint violations. A memory-bounded archive retained at most 200 unique feasible states per search stratum. The final Pareto archive was computed across feasible nonredundant designs by maximizing richness and minimizing changed decoder units, changed codon types, minimum nucleotide substitutions, directional missense cost and GC cost^20–21^. Reported designs are therefore feasible solutions found by the declared search, not certificates of global optimality.

### 2.5 Sensitivity analyses, frozen inputs and reproducibility reporting

The equal-weight arithmetic mean was the principal descriptive scalarization. Sensitivity to objective weighting reused the same three empirical utilities on the fixed 100-million-code AAindex-screened library. Weighted scores

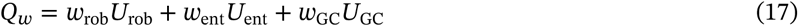

were evaluated over 178 prespecified weight vectors: a grid with increment 0.05 and positive components of at least 0.05, plus named single-objective-emphasis and two-objective stress tests. Arithmetic, geometric and minimum-utility aggregation were compared, and direct three-objective Pareto comparisons used a numerical tolerance of 10^−7^.

Sensitivity to screening design jointly varied the AAindex discoverytail fraction *q* ∈ {0.05, 0.10, 0.20} and three independent seed sets per threshold. Each run repeated rule learning using 90,000,000 training permutations, generated a fresh 10-million-code library by exact rank–unrank sampling and recalculated all three objectives. Because a change in *q* changes the allowed-assignment matrix, these comparisons concern distinct conditional libraries rather than measurements within one common null ensemble.

The revised analysis identifies its three frozen inputs by the following author-reported SHA-256 values: AAin-dex matrix, `33c057d49ab2cd75350050cd804811620c97c59c1e55ba364e03655ad0df4781`; ProteinGym matrix, `05ad196bf47c4ab59cd2ca4f1f36dd2655804983cc68c6fa012dd2cd355c166c`; and MG1655 proteome, `3c548c41721d3adbcc01b4cba1989fafa055e1406422956c5e5cf2ada57116c5`. These identifiers and all computation-derived quantities require the corresponding frozen files and executable workflow for independent verification.

## 3. Results

### 3.1 Genetic-code space decomposes into nested, non-interchangeable structural layers

A genetic code written directly as an arbitrary function from 64 codons to 20 amino acids and one termination signal has a nominal search space of 21^64^. Although this expression conveys the scale of the problem, it conflates biologically distinct questions: why codewords have length three, how 64 codons are distributed among 21 outputs, why synonymous codons form blocks of their observed sizes and shapes, and why those blocks carry their present amino-acid meanings. We therefore represented a genetic code over the nucleotide alphabet Σ={A,C,G,U} as a sequence of nested variables:

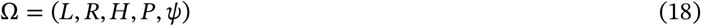

Here, L is the codeword length; H=(h_1_,…,h_21_) is the multiset of degeneracies assigned to the 21 outputs, with Σ_*i*_h_*i*_=4^*L*^; R=max_*i*_h_*i*_ is the maximum degeneracy; P={B_1_,…,B_21_} is an anonymous partition of Σ^*L*^ satisfying |B_*i*_|=h_*i*_; and ψ:P→*A* assigns the anonymous blocks to 20 amino acids and a termination signal. The resulting dependency order is:

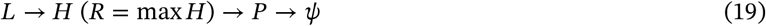

These variables are not independent parameters. Fixing one layer restricts the next layer to a conditional subspace: a single degeneracy profile H can support many codon partitions P, and a single partition can support many semantic assignments ψ. This decomposition separates code capacity, redundancy allocation, synonymous-block geometry and amino-acid semantics into distinct, testable problems.

The decomposition also determines which evidence is meaningful at each layer. Amino-acid properties, codon adjacency, transfer ribonucleic acid (tRNA) decoding and protein sequences are undefined at the L layer, which can therefore be evaluated only through capacity, length cost, information efficiency and the optimal mutation-exposure envelope attainable after partitioning^9,18^. At the H/R layer, codons have not yet been assigned to specific blocks, so only the composition of the 21 block sizes and the downstream partitions permitted by that composition can be compared. Entropy, the Gini coefficient or maximum degeneracy cannot distinguish different geometric arrangements of blocks with identical sizes. At the P layer, the codon members of each block are specified but amino-acid labels remain absent; single-nucleotide adjacency, translational confusion and decoder coverage can therefore be evaluated, whereas amino-acid replacement damage cannot. Amino-acid replacement costs, proteome composition and semantic rewiring become well-defined only at the ψ layer. Thus, different structural features of the genetic code require different randomization spaces, and every result must specify both the layer being tested and the downstream structure held fixed.

At the L layer, we fixed a four-letter alphabet, 21 equally weighted outputs, a uniform codeword length and use of all 4^*L*^ words, without invoking H, P or ψ. The capacity constraint is:

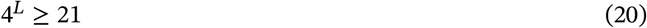

The 16 words available at L=2 cannot cover 21 outputs, whereas L=3 is the first feasible length. This capacity bound establishes only that triplets are sufficiently short; it does not exclude the possibility that longer codewords exploit additional redundancy to obtain greater robustness. We therefore constructed the quaternary Hamming graph on Σ^*L*^ and minimized the 21-way RatioCut over all surjective 21-block partitions^9^. Exact edge-isoperimetric profiles, dynamic programming over block-size compositions and blockwise reachability audits gave the same global minimum for L=3, 4, 5 and 6:

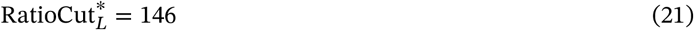

The corresponding optimal macro-averaged robustness was:

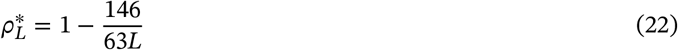

After correction for the number of nucleotide-level mutation opportunities, the effective mutation exposure was:

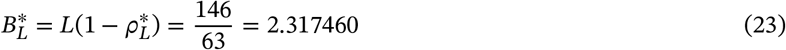

This value remained unchanged from L=3 to L=6. Longer codewords increased the conditional probability that a mutation remained within the original block, but did not reduce optimal output-changing exposure after accounting for the additional mutable nucleotide positions. At the same time, sequence-length cost increased from 1 at L=3 to 2 at L=6 when measured relative to triplets, whereas information efficiency was:

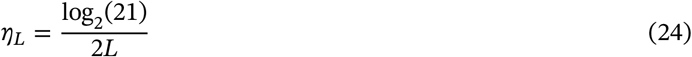

Information efficiency decreased from 0.7321 to 0.3660. Under this fixed-length, full-codebook and equal-weight single-nucleotide model, L=3 therefore achieved the same optimal effective mutation exposure as longer codewords while minimizing nucleotide cost and maximizing information efficiency. Triplets consequently formed a strict Pareto choice within this model. This result applies to the fixed-length total-map branch; sparse codebooks, variable-length codes and systems that require external synchronization occupy different candidate spaces.

After fixing L=3, 64 codons must be distributed among 21 non-empty blocks. If amino-acid identities are ignored and block sizes are sorted in nondecreasing order, every possible H corresponds to a restricted integer partition of 64 into 21 positive parts^10^. Exact enumeration yielded 59,755 anonymous degeneracy profiles. Including the termination block, the standard genetic code (SGC) has the profile:

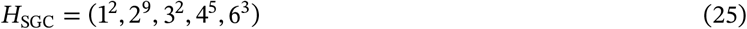

This profile contains two one-codon blocks, nine two-codon blocks, two three-codon blocks, five four-codon blocks and three six-codon blocks, giving R_SGC_=6. Although it lies between uniform and extremely concentrated allocations, that intermediate position did not distinguish it from the 59,755 candidates. For every profile, we calculated anonymous summary descriptors including maximum degeneracy, Shannon entropy, the Gini coefficient, the Herfindahl-Hirschman index and collision probability. Under the pilot descriptor set, 50 noncanonical profiles Pareto-dominated H_SGC_. The canonical profile was therefore neither the most uniform nor the entropy-maximizing or least-concentrated allocation; R=6 is a coarse projection of H rather than an independent explanatory variable.

Natural code variation further exposed the information boundary of H. In Euplotes, reassignment of UGA from termination to Cys expands the Cys block from two to three codons and contracts the Stop block from three to two^24^. The two block sizes are exchanged, leaving the sorted multiset H unchanged even though both P and ψ have changed. Any statistic that depends only on H is blind to this natural recoding event. Consequently, the biological value of H must be evaluated through the downstream partitions, decoders and semantic assignments that it can support. The exact enumeration therefore establishes a limitation rather than deriving the canonical profile from an anonymous balance principle: block-size statistics alone do not explain canonical degeneracy, and additional constraints must enter at the P and decoder-implementation layers.

At the P layer, the question changes from how many codons occupy each block to which codons belong to the same block. This analysis requires three distinct structures: the mutation graph generated by single-nucleotide substitutions, the translational-confusion graph generated by near-cognate tRNA misreading, and the decoding hypergraph in which one decoder may recognize multiple codons^11–14^. Because these structures have different edge weights and biological origins, they cannot be merged a priori into a single unweighted Hamming graph. We first constructed an exactly enumerable local P space from the six canonical family boxes with strict 2+2 splitting at the third codon position^13–14^. Each four-codon box admits three perfect matchings that divide the four third-position nucleotides into two doublets, giving:

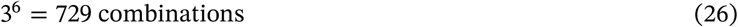

Let transition edges have weight κ and transversion edges have weight 1. The canonical pyrimidine/purine split U/C | A/G has cross-block cut 4 in each box, whereas either alternative pairing has cut 2κ+2. When κ>1, the canonical U/C | A/G split uniquely minimizes mutation flow across synonymous blocks within this local subspace. When κ=1, all three pairings have equal cost and all 729 six-box combinations are tied. The standard doublet orientation at the third codon position is therefore not selected by the premise that the third position is intrinsically less important; its local advantage emerges from the interaction between substitution anisotropy and block geometry. An unweighted Hamming graph cannot select the canonical orientation, whereas unequal transition and transversion weights break the symmetry. Because this calculation covers only six 2+2 family boxes, it provides exact local evidence for a P-layer mechanism rather than a global optimality proof over all partitions of 64 codons.

Having separated L, H and P conceptually, the large-scale analysis fixed L=3, the 20 canonical sense-codon blocks and the positions of the three termination codons, and varied only the bijection ψ between sense blocks and amino-acid labels, matching the fixed-block randomization space used in prior error-minimization analyses^15–16^. Because the termination block was excluded from amino-acid label permutation, the conditional candidate space was exactly:

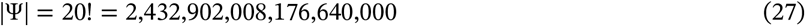

Each candidate in this space is not an arbitrary 64-to-21 function. It is a complete genetic code that preserves the canonical degeneracy composition and synonymous-block geometry while reassigning the meanings of the 20 amino acids. All subsequent large-scale results on rule learning, space contraction, mutational robustness, encoding entropy, local guanine–cytosine (GC) stability and the rank of the SGC therefore answer a strictly conditional question: given that the canonical triplet-block architecture has already formed, does the present amino-acid assignment occupy a favorable region? These results do not constitute a proof over the complete space of L, H and P combinations.

### 3.2 An AAindex recurrent-rule screen contracts the fixed-block assignment space

Conditioning on the canonical synonymous blocks reduced the semantic assignment problem to exactly 20! = 2.43 × 10^18^ permutations, but this remained too large for exhaustive downstream evaluation. The recurrent-rule screen therefore learned local block–amino-acid assignments depleted from the low-cost tail defined by the symmetric AAindex distance. Nested recurrence across three rounds within each of three independent seed groups stabilized the proposal constraints while protecting every canonical assignment.

At the reference threshold *q* = 0.10, the screen yielded 196 recurrent exclusions. Exact subset dynamic programming showed that these rules retained 271,542,412,800 complete mappings, or approximately 2.72 × 10^11^. Rank–unrank sampling then generated 100,000,000 mappings uniformly from this retained space. The reduction is conditional on the declared AAindex matrix and recurrence rule. It is not a classification of biological viability.

Importantly, the screen was independent of the directional ProteinGym objective used downstream. Candidate mappings were filtered using a symmetric physicochemical distance before ProteinGym mutation costs, proteome composition, entropy or GC quantities were evaluated. This separation reduces direct reuse of the downstream mutation score as its own proposal target, although both stages still encode chosen notions of amino-acid similarity.

**Figure 1.**
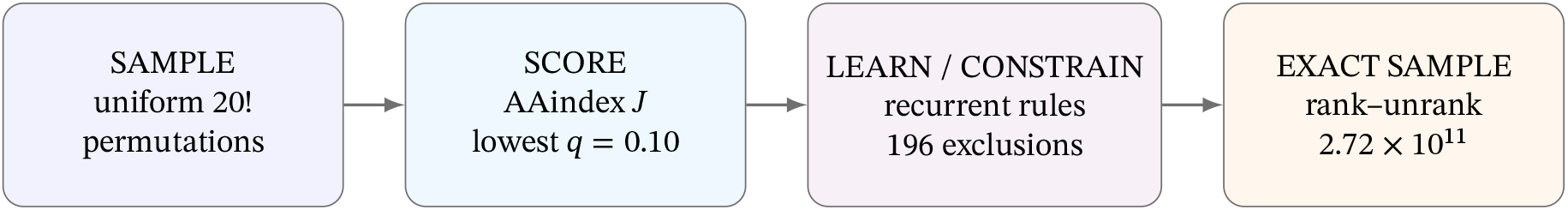
AAindex-based recurrent-rule screening and exact sampling of the fixed-block genetic-code space. Uniform amino-acid label permutations are screened with a symmetric AAindex physicochemical distance, recurrently compiled into an allowed-assignment matrix and sampled by exact rank–unrank. Exclusions are proposal constraints rather than biological impossibility statements.

### 3.3 The SGC occupies a conditional three-objective compromise

We evaluated the standard genetic code (SGC) in the AAindex-screened library using three objectives calculated after candidate generation: directional ProteinGym mutation robustness, MG1655 encoding entropy under uniform synonymous-codon use and local GC stability under the same convention. The SGC ranked in the best 0.235% for directional mutation robustness and 0.0121% for encoding entropy, whereas local GC stability lay in the best 39.3%. The corresponding empirical utilities were 0.998, 1.000 and 0.607. Thus, robustness and conditional encoding capacity were the strongest individual signals, whereas local GC stability limited the composite performance.

Under the equal-weight empirical-rank score *Q*_3_, the SGC ranked 974,903rd among 100,000,000 mappings, placing it in the best 0.9749% of this conditional library. High-*Q*_3_ codes were enriched at smaller label-transposition distance from the SGC. This distance is a combinatorial property of complete block-label assignments and should not be interpreted as a reconstructed evolutionary trajectory.

**Figure 2.**
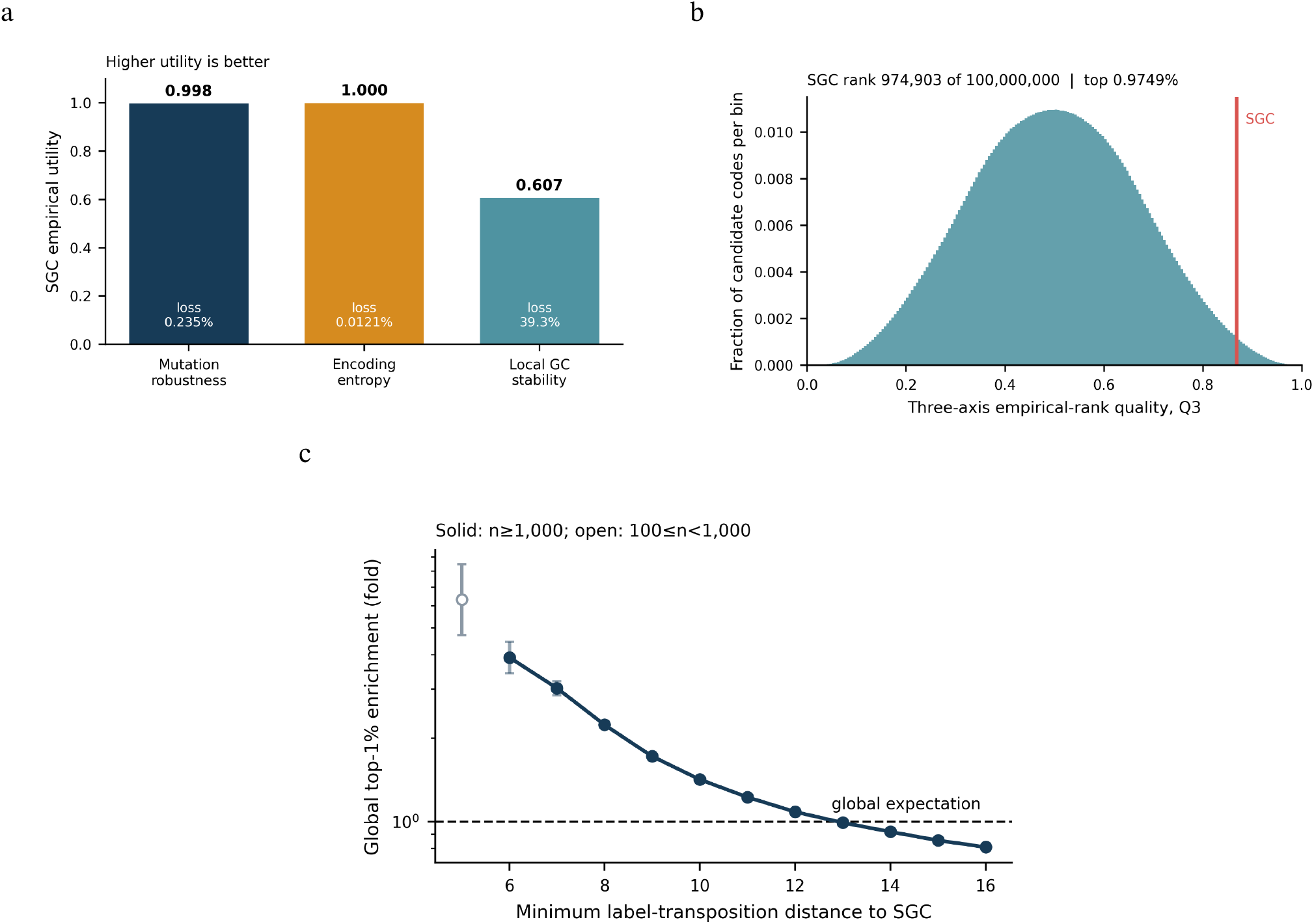
Conditional three-objective position of the SGC in the AAindex-screened fixed-block library. (a) Empirical utilities for directional mutation robustness, encoding entropy and local GC stability. (b) Equal-weight *Q*_3_ distribution; the red line marks the SGC (rank 974,903 of 100,000,000; top 0.9749%). (c) Top-1% *Q*_3_ enrichment by minimum block-label transposition distance.

### 3.4 Accessible replacement diversity exposes a robustness–explorability trade-off

Accessible replacement diversity was added as a fourth objective. It measures the directional ProteinGym separation of two independently sampled sense-missense targets accessible from the same source amino acid under the declared codon and mutation sampling process. This is an evolvability-like design axis, not an estimate that an accessible mutation is beneficial.

With this fourth objective, the SGC retained high utilities for directional robustness (0.998) and entropy (1.000), but lower utilities for local GC stability (0.607) and accessible replacement diversity (0.381). The equal-weight score *Q*_4_ placed the SGC 1,801,376th of 100,000,000 mappings, or in the best 1.801% of the AAindex-screened conditional library. The shift from *Q*_3_ demonstrates that apparent extremeness depends on the declared objective set.

The decoder-unit search remains a constrained design exercise rather than evidence of cellular implementation. Its output is not used to establish the principal conditional ranking. A compatible tRNA, aminoacyl-tRNA synthetase, modification, ribosome and genome-wide recoding system would require separate molecular design and validation.

### 3.5 Sensitivity analyses delimit the conditional result

Across 178 tested weight vectors, the SGC was within the best 1% for 86 (48.3%), the best 5% for 131 (73.6%) and the best 10% for 149 (83.7%). Its ranking was strongest when entropy received substantial weight and weakest under GC-dominated weights; the tested vector (0.05, 0.05, 0.90) placed it in the best 33.876%. Its percentiles from best were 0.9749%, 1.4078% and 6.1945% under arithmetic-mean, geometric-mean and minimum-utility aggregation, respectively. One of the 100,000,000 mappings dominated the SGC on all three utilities, so the SGC was not exactly on the observed three-objective Pareto frontier.

Independent AAindex rule learning was stable across *q* ∈ {0.05, 0.10, 0.20}. Mean equal-weight SGC percentiles in new 10-million-code libraries were 0.874885%, 0.977903% and 0.939007%, with within-threshold rule-set Jaccard similarities of 0.9966, 1.0000 and 0.9966. These checks support robustness to the declared screen and scalarizations, but each threshold generates a distinct conditional library and does not establish a global rank over all 20! mappings.

**Figure 3.**
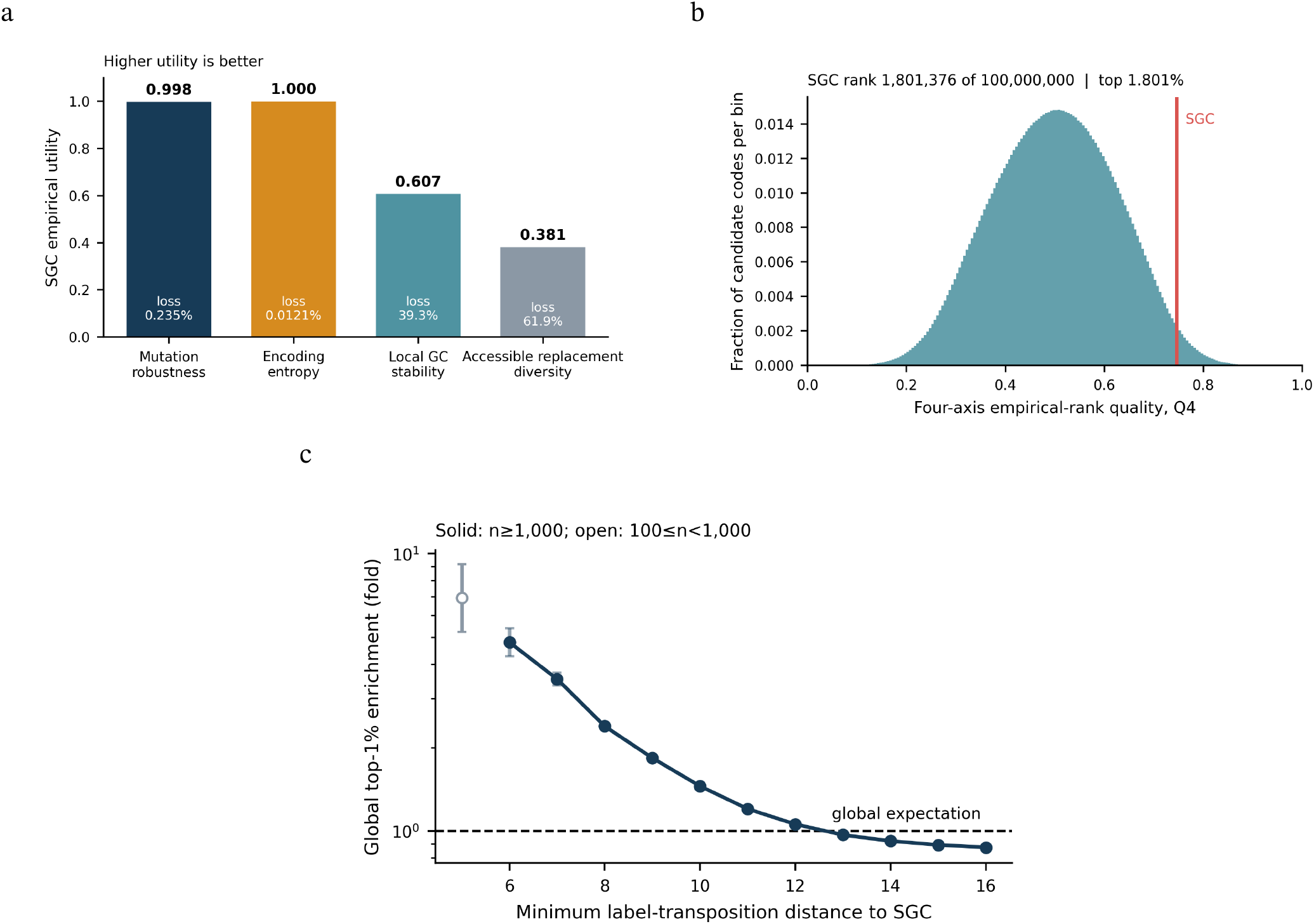
Accessible replacement diversity exposes a robustness–explorability trade-off without eliminating the SGC’s conditional multi-objective signal. (a) Four empirical utilities. (b) Equal-weight *Q*_4_ distribution; the red line marks the SGC (rank 1,801,376 of 100,000,000; top 1.801%). (c) Enrichment of global top-1% *Q*_4_ codes by label-transposition distance.

## 4 Discussion and Conclusion

This study treats the canonical genetic code as a hierarchy of conditional design problems rather than an unrestricted mapping from 64 codons to 21 outputs. The structural analyses concern distinct codeword, degeneracy and block-partition layers. The large-scale semantic result begins only after the canonical triplet sense-block architecture and Stop positions have been fixed. Consequently, evidence at one layer cannot by itself explain choices made at another.

Under the fixed-length, full-codebook model, triplets form a strict Pareto choice: *L* = 3 is the shortest feasible code-word length, whereas longer words from *L* = 4 to *L* = 6 do not reduce effective mutation exposure after their additional mutable positions are counted. The degeneracy analysis gives an equally important negative result. Among 59,755 anonymous block-size profiles, 50 noncanonical profiles Pareto-dominate the canonical profile under the pilot descriptors. Maximum degeneracy *R* = 6 is therefore a constraint on the search space, not a sufficient explanation of the canonical allocation. Both results remain conditional on their stated graph, capacity and descriptor assumptions.

The strongest positive result appears after the canonical triplet-block architecture is fixed. A symmetric AAindex screen, independent of the downstream directional ProteinGym score, retained 2.72 × 10^11^ semantic assignments. In the resulting 100-million-code library, the SGC had strong but nonuniform performance: top 0.235% for directional mutation robustness, top 0.0121% for conditional encoding entropy and top 39.3% for local GC stability. Its equal-weight composite ranks were top 0.9749% under *Q*_3_ and top 1.801% after accessible replacement diversity was added in *Q*_4_. The SGC is therefore better described as an unusual compromise than as the optimizer of any single objective.

Sensitivity analysis limits the strength of that conclusion. The SGC remained within the best 10% for 83.7% of the tested weight vectors, but fell to the best 33.876% under the most GC-dominated tested vector and to 6.1945% under minimum-utility aggregation. One sampled mapping also dominated it on all three principal utilities. The evidence thus supports broad conditional performance across many declared priorities, not invariance to weighting, exact Pareto optimality or a universal rank.

Several abstractions remain biologically important. Uniform synonymous-codon sampling does not reproduce native codon bias or tRNA demand; amino-acid occurrence counts do not represent expression-weighted cellular burden; the unweighted block graph does not reproduce the MG1655 mutation spectrum or translational misreading; and local GC deviation is not a model of mRNA folding, initiation or transcript stability. The project-specific ProteinGym matrix also aggregates heterogeneous measurements and cannot capture residue-specific structural or functional effects without additional modelling. These limitations do not invalidate the conditional comparison, but they prevent its scores from being interpreted as cellular fitness.

The decoder-unit analysis should likewise be read as a coarse candidate-design exercise. It does not show that a compatible set of tRNAs, aminoacyl-tRNA synthetases, RNA modifications, ribosomal interactions and genome-wide recoding operations can implement a proposed mapping in cells. Future work should replace the current uniform and pooled assumptions with native MG1655 codon usage, measured mutation spectra, tRNA abundance, wobble pairing, mistranslation matrices, expression-weighted proteomes and protein-level structural tests. Overall, the SGC is a conditionally unusual multi-objective compromise under the fixed architecture and specified objectives examined here, not a demonstrated global optimum, historical inevitability or validated synthetic-cell design.

## 5 Data and Code Availability

The revised analysis specifies frozen AAindex and directional ProteinGym matrices, an MG1655 proteome file, objective weights, pipeline modules, pseudocode, tests and source tables for Figure 4. The three input identifiers are reported in Methods. These files and the executable workflow were not distributed with this manuscript version; the computation-derived values and hashes therefore remain author-reported pending independent reproduction. Before public release, the complete package should be deposited in a persistent repository with versioned source code, environment information and a command that regenerates every reported table and figure. The 100-million-code HDF5 files need not be redistributed if the documented workflow deterministically regenerates them from the frozen inputs and seeds.

**Figure 4.**
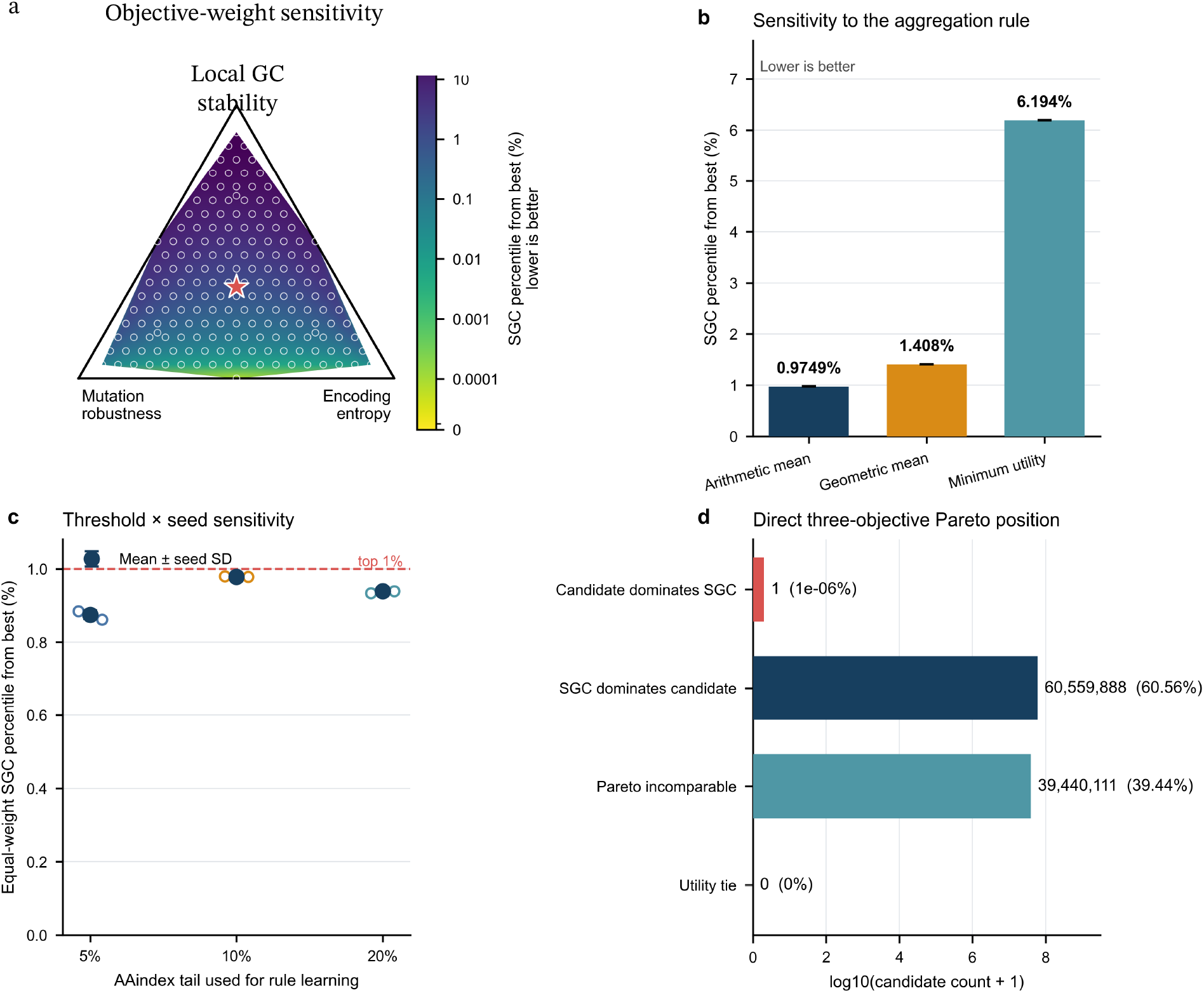
Sensitivity of the conditional three-objective position of the SGC. (a) Tested objective weights. (b) Al-ternative aggregation rules. (c) AAindex discovery-tail and seed sensitivity. (d) Direct Pareto comparison. Lower percentiles are better; the displayed threshold libraries are not one common null ensemble.

## Acknowledgements

This work was developed during the 2026 PEBBLE BioFusion Workshop held at Westlake University in Hangzhou, China, from July 24 to August 4, 2026. All authors gratefully acknowledge the speakers for their inspiring lectures and discussions and the teaching assistants for their generous guidance and support throughout the workshop. We especially thank Dr. Fangzhou Xiao, the workshop organizer, whose enthusiasm, energy, dedication and generous assistance strongly supported the development of this project and enriched the workshop experience.

